# Hippocampal Place-like Signal in Latent Space

**DOI:** 10.1101/2022.07.15.499827

**Authors:** Matthew Schafer, Philip Kamilar-Britt, Vyoma Sahani, Keren Bachi, Daniela Schiller

## Abstract

During navigation, the hippocampus represents physical places like coordinates on a map; similar location-like signals have been seen in sensory and concept spaces. It is unclear just how general this hippocampal place code is, however: does it map places in wholly non-perceivable spaces, without locations being instructed or reinforced and during navigation-like behavior? To search for such a signal, we imaged participants’ brains while they played a naturalistic, narrative-based social interaction game, and modeled their relationships as a kind of navigation through social space. Two independent samples showed hippocampal place-like signals in both region-based and whole-brain representational similarity analyses, as well as decoding and average pattern similarity analyses; the effects were not explained by other measures of the behavior or task information. These results are the first demonstration of complete domain generality in hippocampal place representation.

**One-Sentence Summary:** hippocampal place-like signal in non-perceivable and unreinforced space during naturalistic navigational behavior.

**Significance statement:** The hippocampus is a brain structure known to encode maps of physical spaces; this study shows that it also maps fully abstract, latent and uninstructed spaces. People played a naturalistic social interaction game while their brains were scanned. Hippocampal brain activity correlated with the fictional characters’ locations in an abstract social space framed by axes of affiliation and power, despite the participants never being exposed to a perceivable spatial representation. This mapping was present across multiple analyses and two samples, demonstrating that the brain system responsible for spatial mapping maps our social interactions too.

## INTRODUCTION

Mapping is an efficient way to organize spatial information and aid navigation. Neural maps may be encoded in the hippocampus, with cells that track the physical locations of oneself (O’Keefe and Nadel, 1978), others (Omer et al., 2018), objects (Manns and Eichenbaum, 2009) and rewards (Gauthier and Tank, 2018). Hippocampus place mapping extends beyond physical environments: similar effects are seen for moments in time (Eichenbaum, 2014) and sensory experiences (Aronov, Dmitry, Nevers, Rhin, Tank, 2017), often in the same cells that track physical locations. Even concept space is mapped similarly. In one study, human participants were taught to associate different stimuli with specific locations in a visual feature space (Theves et al., 2019). After the learning phase, hippocampus activity showed a coordinate-like pattern: like a map, the similarity between representations of different locations reflected the relative distances between them (i.e., closer locations were represented more similarly). Altogether, these findings suggest that the hippocampus maps relationships regardless of the domain (i.e., is domain general) (Behrens et al., 2018; Bellmund et al., 2018; Schafer and Schiller, 2018). Here, we sought to test for complete domain generality in the hippocampal place code by examining an extreme case.

Domain general place codes should abide by several key principles. The first is the ability to map fully abstract information. To date, place mapping has been shown in spaces anchored by perceivable dimensions (e.g., variations in an object’s visual features). However, any behaviorally relevant dimension should be mappable, even those that are not directly perceivable and can only be inferred through their influence on other observable features (i.e., total abstraction). Second, learning phases where the participant memorizes the map shouldn’t be necessary: mapping should happen even when the dimensions are uninstructed and the locations are unreinforced. Third, place representations should be apparent during naturalistic navigation-like behavior: they should emerge spontaneously and implicitly when exploring natural variation, not only when participants are explicitly required to organize information into a map-like format. In sum, if the hippocampus place code is completely domain general, place representations will emerge during the unsupervised and naturalistic navigation of non-perceivable spaces – reflecting a fully latent and relational mapping of abstract information.

To test for domain general hippocampal place-like representations, we used social interaction. Adaptively interacting with others is akin to an act of navigation: it may depend upon mapping and using people’s locations along latent social dimensions to guide social decision-making, and updating these locations as more is learned and relationships change (Schafer and Schiller, 2018; Tavares et al., 2015). Affiliation (e.g., bonding, friendship) and power (e.g., dominance, competence) are two dimensions that may anchor an abstract social space: people’s social place on affiliation and power is important to many social processes, including trait attribution from faces (Todorov et al., 2008) and stereotyping (Fiske, 2012) and resource sharing and mating in non-human primates (Feldblum et al., 2021; Samuni et al., 2018). Importantly, these dimensions cannot be directly perceived and must be inferred, such as through their effects on social interactions. In this study, we ask whether the hippocampus represents others as dynamic locations (i.e., latent place coding) in a social power and affiliation space during naturalistic social interactions.

If the hippocampus maps people as places onto latent social dimensions, we predict the following. First, hippocampus representational similarity between social places will vary as a function of their distance in latent space: close locations will be represented more similarly than far apart locations – as if coordinates in a map. Second, because social relationships are ongoing and change over time, social place representations should dynamically update as relationships evolve. Third, mapping should be independent of individuating social information, like a person’s identity (though conjunctive representations of place and person may exist in the hippocampus, like place and object conjunctions in physical space, e.g., Komorowski et al., 2009). Fourth, other functional properties of place codes may be present: for example, a posterior-to-anterior hippocampal gradient for increasing place field size (Brunec et al., 2018). Fourth, we should be able to detect the social place representations as people navigate their own social spaces during naturalistic interactions without an explicit requirement to express the characters as locations. And, fifth, the representations should not require any instructions or explicit reinforcement about the underlying dimensions or locations.

We collected two independent samples (n=18 and n=32) on a “choose-your-own-adventure” game that is both social and naturalistic: participants made decisions in a variety of simulated social interactions with fictional characters over the course of a narrative. These decisions implicitly updated the characters’ coordinates on (non-perceivable) social dimensions of affiliation and power, producing a sequence of latent two-dimensional locations across time for each character – their trajectories through social space (**fig. 1**). There were no instructions or learning phase to pre-define the dimensions or locations, or task requirements to locate the characters spatially. Evidence for place mapping under these extreme conditions would be evidence for a true domain general process during navigation-like behavior.

**Fig 1.**
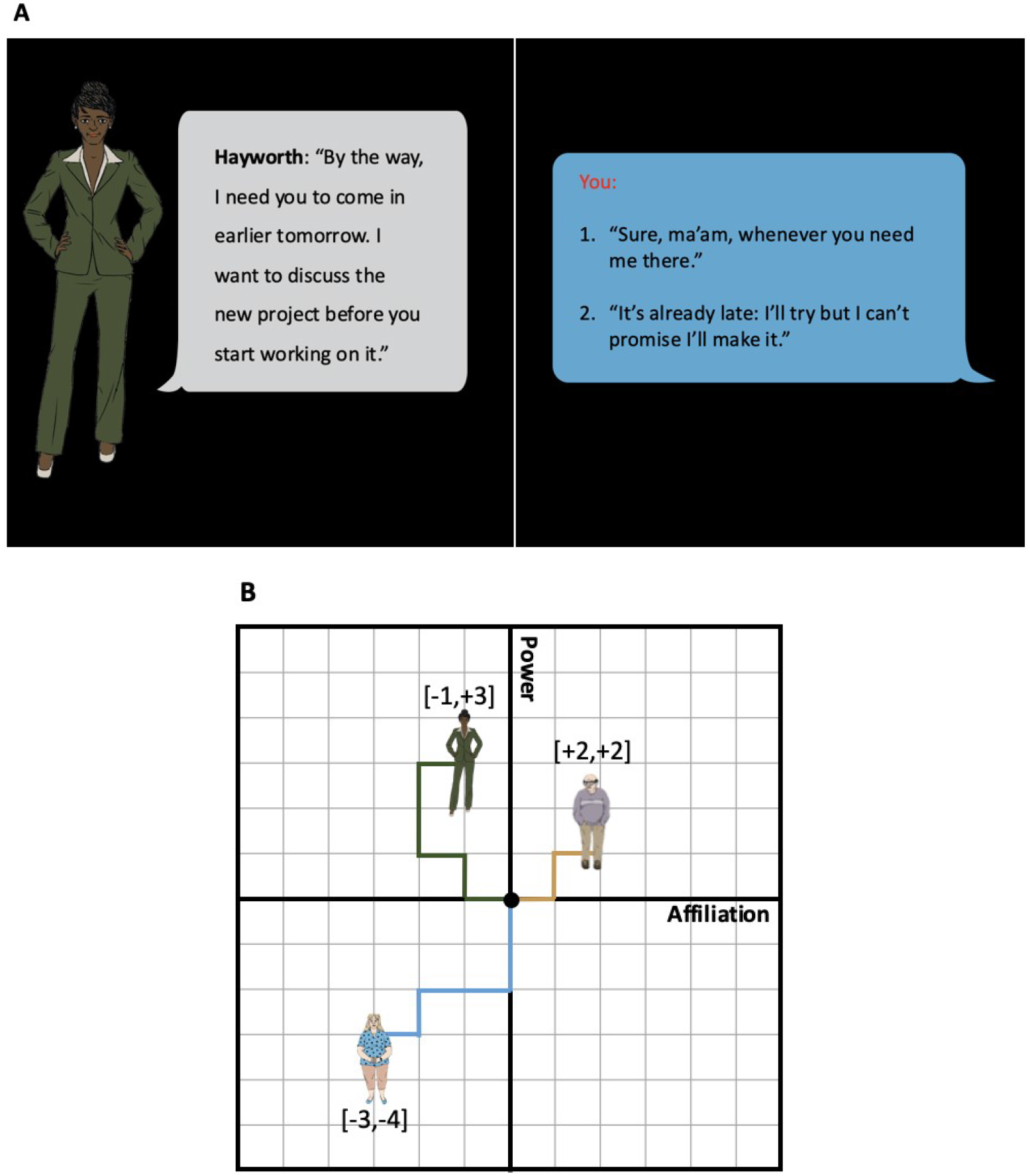
Social navigation task and latent place modeling. (A) An example of a power decision. Participants read text that describes the narrative and on decision trials choose between two options. Based on their choice, the character moves −1 or +1 along the active dimension (affiliation or power). (B) Participants’ decisions were modeled as a sequence of affiliation and power coordinates (i.e., a trajectory). Trajectories are color-coded by character.

## METHODS

### Participants

The initial sample was collected for a previous study (Tavares et al., 2015), and included 21 participants. 18 (7 female) were included after excluding 1 for excessive head motion in the scanner (multiple translations > 1 voxel) and 2 for dicom file corruption during archiving. The validation sample included 32 of 39 participants (17 female) after excluding 2 participants for poor post-task memory (less than chance level) and 5 participants for motion. The two samples did not differ on participant sex (χ^2^=0.452, *p*=0.501) but were significantly different on age (in years; Initial sample: *M*=29.44, *SD*=3.24; Validation sample: *M*=45.17, *SD*=9.8; Wilcoxon rank-sum test: W=518.0, *p*=3.5e-6).

The Institutional Review Board of the Icahn School of Medicine at Mount Sinai approved the experimental protocols for both samples. All participants provided written informed consent and were compensated for their participation (for more information about the samples, see **Sample details** in the supplemental material).

### Social Navigation Task

The social navigation task is a narrative-based social interaction game. Pre-task instructions were minimal: participants were instructed to not overthink their responses and just behave as they would naturally. At the start of the narrative, the participant is told they have just moved to a new town and need to find a job and a place to live. They interact with fictional characters over the game and occasionally have to make decisions in those interactions that could alter their relationships (**fig. 1A**). Unbeknownst to the participants, the slides were the same no matter the decisions the participant made; the post-decision slides were written to have narrative continuity regardless of the specific decisions. This ensured that all participants were exposed to the same text (though there was some counterbalancing of the character’s gender and race across participants, see **Social Navigation Task details** in the supplement). Participants had a button response box to select between two options. The task lasted approximately 26 minutes.

Unbeknownst to the participants, the decision trials were categorized as either affiliation or power decisions. Each decision consists of a choice between two options that move the character in either the negative (−1) or positive (+1) direction along the current dimension. There are 5 characters with 6 affiliation and 6 power decisions each, for a total of 60 decisions. Affiliation decisions were operationalized as decisions whether to share physical touch, physical space, or information (e.g., to share their thoughts on a topic). Power decisions were defined as decisions to submit to or issue a directive/command, or otherwise exert or give control. These task dimensions were designed to be orthogonal and do not predict one another in the participants’ behavior (see **Affiliation and power dimensions are orthogonal in behavior** in the supplemental material); thus they are modeled as the orthogonal axes of a Euclidean plane. There was also a neutral character, with 3 neutral decisions that did not alter their social position, bringing the overall number of decisions to 63.

Each character started the task at the neutral origin with neutral affiliation and power (0,0). With each decision, that character’s (latent) coordinates were (implicitly) updated in the positive or negative direction along the current dimension, based on the option chosen by the participant (i.e., each decision moved the character −/+ 1 arbitrary unit along the given dimension). Thus, at any given point in the task, the characters’ 2D coordinates were the cumulative sums of the participant’s affiliation and power decisions in those specific relationships (**fig. 1B**).

After the task, participants were asked 5 memory questions about each of the 6 characters (5 experimental characters plus 1 control character), with 5 options available. Above chance memory for the 5 experimental characters was required for inclusion in this study.

#### Behavioral uniformity test

To test for a social place effect, social places should vary across participants. We used the Rayleigh test of uniformity to test whether the angles from the origin (0, 0) to the average social places (mean of the affiliation and power decisions) significantly differed from the null hypothesis of a uniform circular distribution. If the participants exhibited independent behavior, there would be a uniform distribution of the 2D social angles and the null hypothesis (of circular uniformity around the origin) would fail to be rejected. The angular mean and standard deviation are used to summarize participants’ average 2D social angles.

#### Post-task character placement

Participants may form a mental representation of the characters’ 2D social places. To probe this, after the task we asked participants (in the validation sample) to complete a subjective character placement task. They were instructed to drag-and-drop colored dots representing each of the 6 characters into a 2D affiliation and power space (explained to the participant), according to the participant’s perceptions of their relationships from the task. The characters were to be placed in this 2D space *relative* to the participant; the participant’s theoretical “point-of-view” (see **Participant’s theoretical point-of-view** in the supplement) was represented by a red dot at the far right end of the affiliation axis and directly in the middle of the power axis.

To test whether participants’ subjective representations of the characters related to the way we modeled the characters’ locations, we asked whether the characters’ locations in this placement task were closer to their locations in the social navigation task than would be expected by chance. For each participant, the average Euclidean distance between each character’s task and perception locations was calculated. 1000 randomly generated sets of character locations were generated (from a uniform 2D distribution, where each 2D location had an equal probability of being selected), and the average of their distances was also calculated. Then the participants’ averaged empirical distances and averaged random distances were compared with a right-tailed paired t-test for our prediction that the true distances would be smaller than chance.

### fMRI acquisition and preprocessing details

Both samples’ data were acquired on 3 Tesla (3T) scanners and preprocessed with a standard pipeline: images were slice-time and motion corrected, unwarped, segmented and normalized. The functional images were left unsmoothed for multivoxel pattern analyses. The images were smoothed with a 6mm Full Width at Half-Maximum (FWHM) Gaussian kernel for the univariate parametric modulation analyses. See the **Image acquisition** and **Image preprocessing** sections in the supplement for more details.

### General Linear Modeling for multivoxel pattern analyses

For multivoxel pattern analyses, we regressed the warped functional images onto a design matrix with a regressor per decision trial, a regressor for the narrative trials and regressors of no interest for the motion parameters. Voxels whose average signal exceeded 50% of the global signal were modeled. The beta estimates from these General Linear Models (GLMs) were used for subsequent pattern analyses, often from *priori* regions of interest (ROIs). To make comparison between the samples easier (e.g., so searchlight analyses cover the same number of voxels and volume across the samples), the initial sample’s images were resliced to match the spatial dimensions of the validation sample. The **Trial-wise general linear modeling** and **Region of interest definitions** sections in the supplement have more details.

### Representational similarity regression analyses

Our main hypothesis was that the hippocampus represents social places: hippocampal patterns for nearby social places should be more similar than the patterns for farther apart ones, reflecting a maplike code where people are represented as locations along social dimensions. We tested this prediction with a representational similarity analysis (Kriegeskorte et al., 2008) where we regressed (with a Huber estimator) hippocampal pattern dissimilarities onto place distances and covariates of no interest (**fig. 2D**). For each participant, we calculated the correlation distances (1 – Pearson’s r) between their trial-wise hippocampal patterns and the Euclidean distances between their trial-wise 2D social coordinates; control behavioral models were calculated similarly (see **Control distance matrices** in the supplement). Additional distance matrices were used to account for motor and action-selection processes (reaction time and button press), character identity and familiarity, the narrative structure (slide and scene changes) and time (i.e., temporal pattern drift). Controlling for these effects is essential to isolating the effect of social place itself. See **Representational similarity regression details** in the supplement for more information.

**Fig 2.**
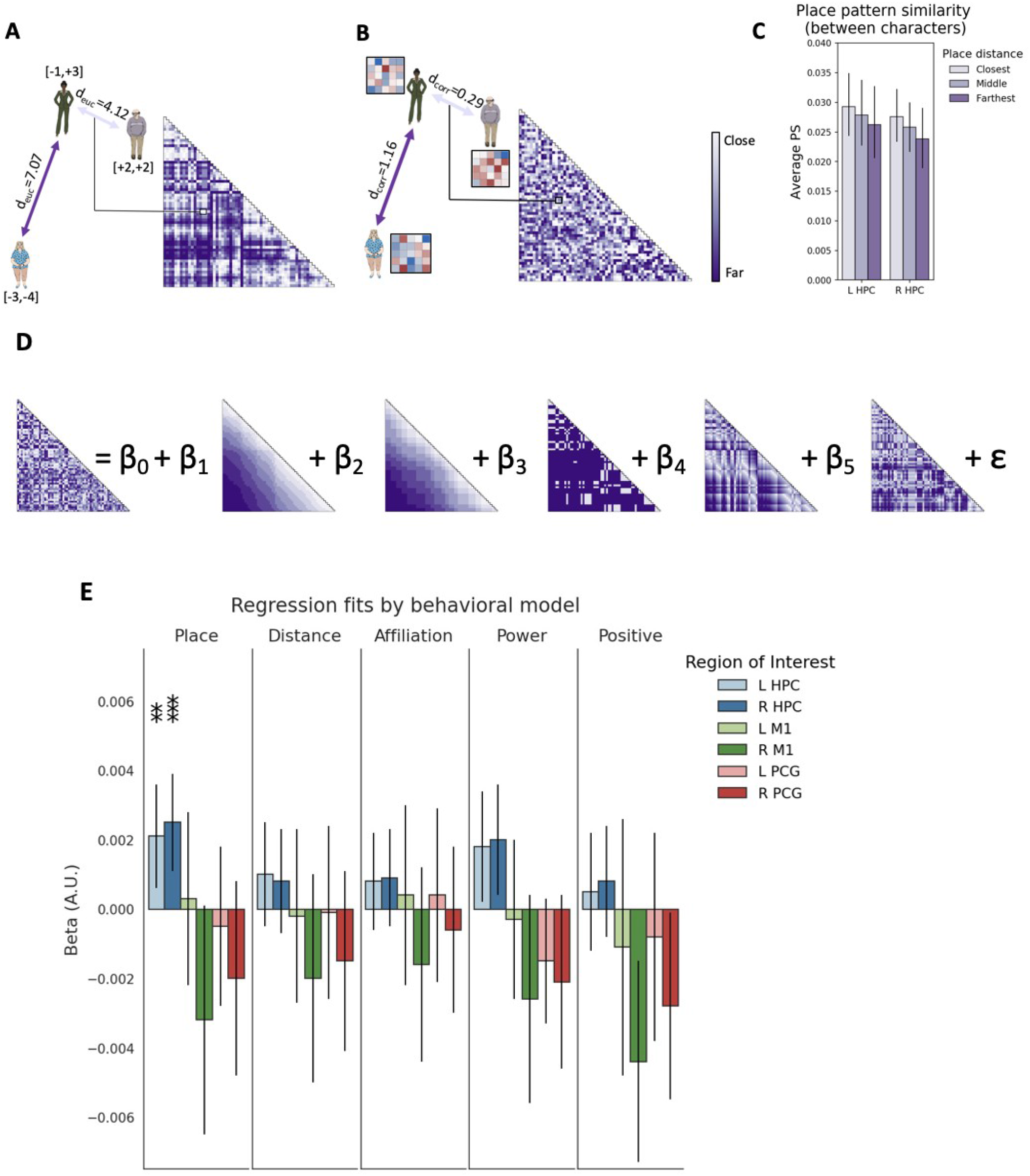
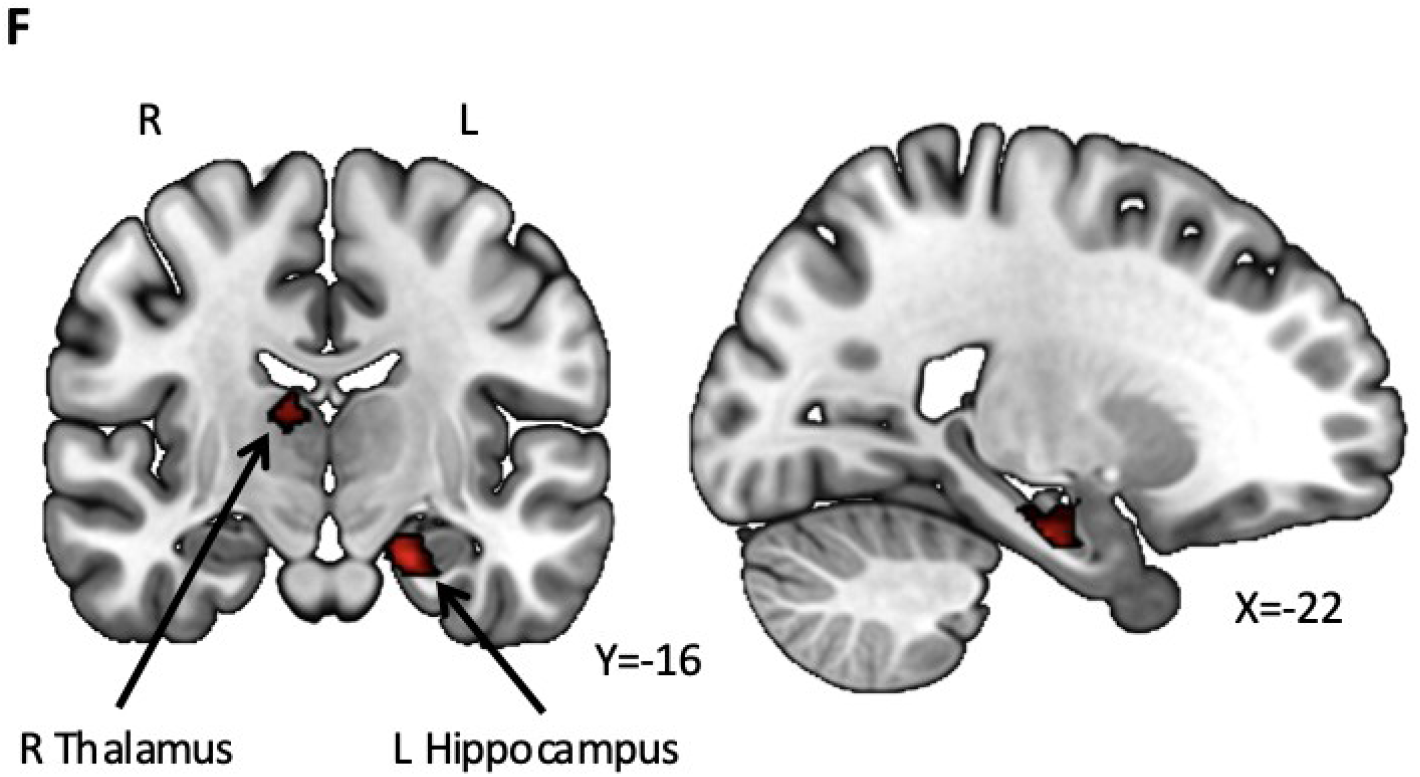
Representational similarity regression shows significant place effects in the hippocampus. (A) For each participant, the pairwise Euclidean distances between each trial’s coordinates were captured in a distance matrix, where each cell is the distance between different trials’ coordinates (small to large distances are indicated with light to dark purple). The distance matrix represents our main prediction: if social places are represented in a map-like format, nearby places should have more similar neural patterns compared to more distant places. Matrices are digitized for visualization purposes. (B) This prediction was compared to the observed neural data for the same participant: patterns of activity from a given region of interest from each trial were extracted and compared in a correlation distance matrix (darker purple means more distance). Matrices are digitized for visualization purposes. (C) The multi-voxel hippocampal pattern similarity (Pearson’s r) between trials increases as the place distance between them decreases. Only between-character trials were used to correct for the influence of within-character trials. (D) A schematic of the multiple representational similarity regression (one participant’s data shown). Predictors from the left are dissimilarity matrices for place distance slide number with the weight to estimate, B_5_, which varies by participant, as well as slide number B_1_, scene number B_2_, character identity B_3_ and familiarity B_4_. Reaction time, button press, and temporal drift dissimilarity predictors, not shown here, were also included as regressors of no interest. Matrices are digitized for visualization purposes. (E) The mean parameter estimates across participants, for the hippocampus (HPC), primary motor cortex (M1) and precentral gyrus (PCG). The results of right-tailed t-tests against chance with 95% confidence intervals are shown; p-values are FWER corrected (Bonferroni) for 2 comparisons (left and right hemisphere) and thresholded: *<0.01, **<0.005, ***<0.001. The place model provided a better fit across the samples for the hippocampus patterns than control models or control regions; there were no differences between samples so their combined effect is shown. (F) Whole-brain searchlight results: clusters in the left anterior hippocampus and the right thalamus survive TFCE correction at *p*_FWER_<0.05. The negative log of p-values are shown.

#### Region of interest (ROI) analyses

Regression analyses were first performed in ROIs: the hippocampus (HPC), our *a priori* region-of-interest, along with control regions of the primary motor cortex (M1) and precentral gyrus (PCG). These regions are likely to be engaged during the task but unlikely to have place cells. The beta estimates from models fit to these regions were computed for all participants and then tested for statistical significance with ANOVA (with partial eta-squared [pes] as the effect size estimate) and follow up t-tests (with Cohen’s d [d] for effect size). 1-tailed 1-sample t-tests were used for directional hypotheses (e.g., right-tailed for the hypothesis that greater distance between places relates to greater dissimilarity between hippocampal neural patterns); 2-sided 95% confidence intervals are reported to give plausible estimate ranges.

Tracking social place may also be related to social memory: we predicted that larger social place beta estimates (possibly reflecting better neural representations of the characters in social space) should positively correlate with post-task memory. Ordinary least squares (OLS) regression was used in the validation sample only; for the initial sample, we only know that the participants performed above chance on their post-task memory – justifying their inclusion in the study but not in this analysis.

#### Whole-brain searchlight analysis

We complemented the ROI regression analyses with a whole-brain searchlight analysis to search for place effects across the brain in an unbiased manner. The regression was computed for every voxel in the intersection of the participants’ GLM-level masks that also had at least 50% probability of being gray matter. For each included voxel, the regression was computed within a spherical searchlight volume (3 voxel radius) and the place parameter estimate was stored. The searchlight sphere covers the same brain volume for all participants: the lower spatial resolution initial sample’s images were resampled to match the higher spatial resolution validation sample images to ensure maximum comparability.

Each participant’s searchlight beta map was spatially smoothed to minimize the effect of inter-subject variability in the spatial localization of the social place effect. We used a small Gaussian smoothing kernel (2.1mm at FWHM, based on a single voxel) because: 1) we wanted to preserve the voxel-level inference, 2) we expected the spatial extent of the effect to be small to moderately sized and 3) the searchlight process is already a kind of spatial smoothing that pools information across voxels. 1-sample right-tailed t-tests were then performed across all participants at each voxel. The resulting t-map was corrected for multiple comparisons with threshold-free cluster enhancement with 5,000 permutations (TFCE; Smith and Nichols, 2009) for family-wise error rate (FWER) control, and thresholded at *p*_TFCE_<0.05. TFCE provides voxel-level correction that is sensitive to the spatial extent of a pattern, without some of the drawbacks of cluster-extent correction methods (e.g., arbitrary cluster-forming thresholds).

### Place pattern similarity analysis

Seeking converging evidence, we tested our main hypothesis again with an approach that made minimal assumptions: we compared the average pattern similarity for decision trials with close versus far social places. We excluded trial pairs where both were from the same character: trials within character are more likely to have both nearby locations (locations are from the same trajectory) as well as high pattern similarity (e.g., shared identity, closeness in time). If we see the hypothesized effect despite excluding within-character trial pairs we will be more confident in interpreting it as a place effect.

Like for the linear and logistic (decoding) regression analyses, we predicted that hippocampal pattern similarity would be greatest for trial pairs with the closest social places. To test this, for each participant we calculated the hippocampal pattern similarities (Pearson’s r) between all trial pairs and then averaged them together based on their social place distances (Euclidean). We binned trials into 3 bins of increasing place distances, balancing the number of trials as closely as possible. We tested the pattern similarity across the bins with a mixed effects model: we modeled average pattern similarity as a function of the fixed effects of social place distance (bins: close, medium, far), hemisphere and sample, and a random effect of participant. The Likelihood Ratio Test was used to test for statistical significance: the model fit was compared to a null model that did not include the distance predictor. To directly compare the effect in the hippocampus and control regions, we used right-tailed paired t-tests to compare average pattern similarity differences between the closest (bin 1) and farthest (bin 3) social place trials, predicting that the hippocampus will show the larger effect.

### Decoding character confusability analysis

Following the regression analysis, we sought converging evidence for a place effect with a separate analysis. We reasoned that if social place signals are in the hippocampus, then models trained to decode character identity from hippocampal patterns would show place-like effects: models trained only on the characters’ identities may incidentally learn social place information and be more likely to “confuse” characters closer in the social space. To test this, a two-step analysis was used. First, for each participant, we trained cross-validated logistic regression models to decode the currently active character from hippocampal patterns (using the 5 experimental characters). To test model performance, we compared the participant-level prediction accuracies against chance (20%) using a 1-sample right-tailed t-test. Second, we compared the trial-wise decoder probabilities for the four non-correct characters based on their end-of-task social place distance to the correct character: the 2 characters closest to the active character should have higher probabilities than the 2 farthest characters. We analyzed the participant-level confusion matrices excluding the correct character (dropping the diagonal, leaving a 5 x 4 matrix). For each trial, we averaged the probabilities for the 2 closest characters and for the 2 farthest characters, and divided them by the total probability assigned to the 4 non-correct characters (producing trial-wise probabilities that sum to 100%). We then averaged these across trials. ANOVA and follow-up right-tailed paired t-tests were used to statistically compare the confusion probabilities for the close and far characters. See **Fig. 3A-B** and **Decoding probability analysis details** in the supplement for more information.

**Fig 3.**
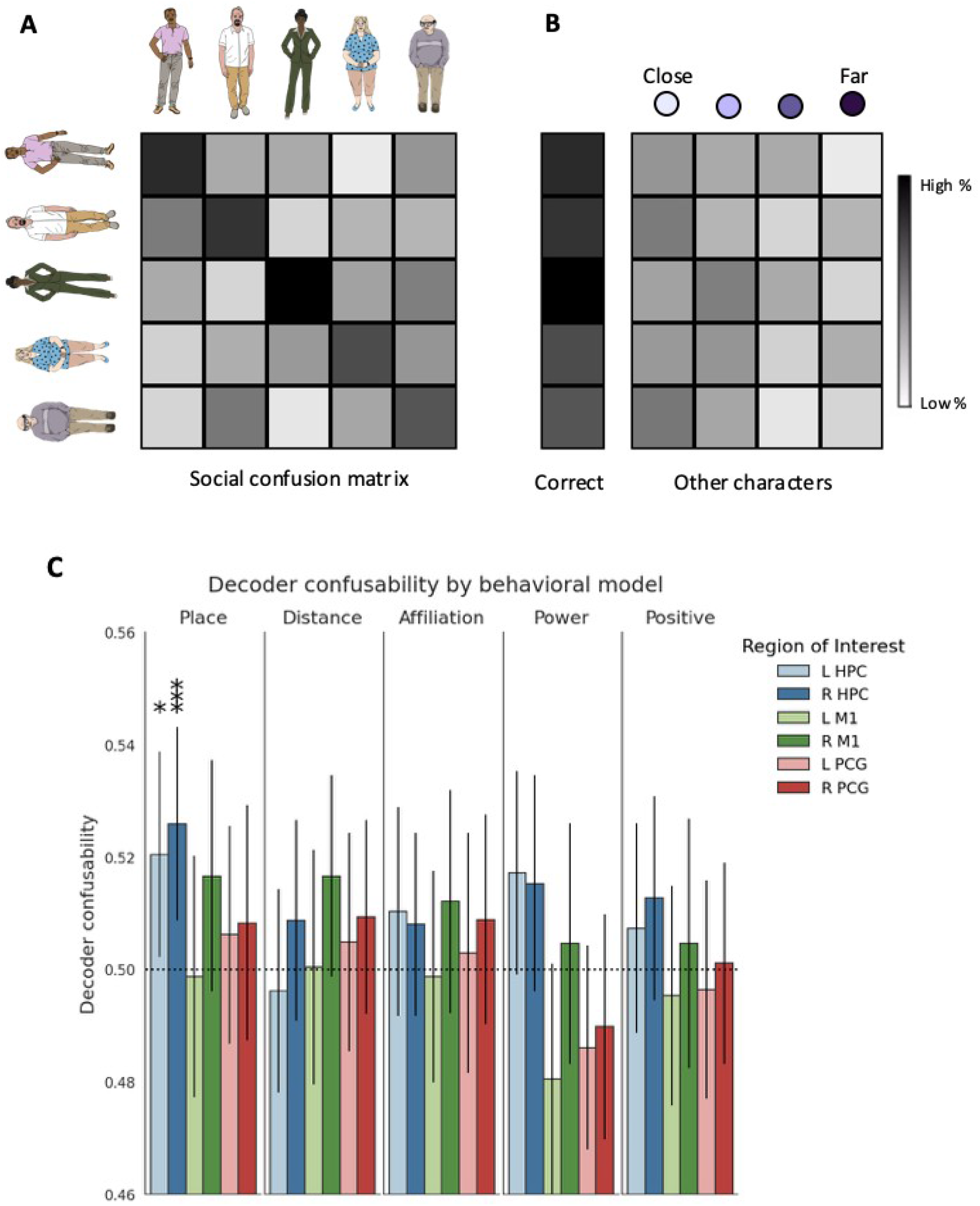
Decoder character confusability varies with place distance in the hippocampus. A) The character decoders produce probability estimates that can be organized into a confusion matrix: the rows indicate the correct characters and the columns indicate the predicted characters. The on-diagonal estimates are the decoded probabilities for the correct characters. Everything off-diagonal is the probability estimated for an incorrect character. B) The diagonal elements (correct) are analyzed to get the accuracy of the decoder. The off diagonal elements (other characters) are examined for their relationship to the place distance: the closest two characters are contrasted to the farthest two characters. C) The results from the two samples are combined, for the left (L) and right (R) hippocampus (HPC), primary motor cortex (M1) and precentral gyrus (PCG). The bars show the decoding probability for the two closest characters, as a proportion of the non-correct probability estimate (i.e., with the correct characters’ estimates removed). The dotted line is chance (50%). The results of right-tailed t-tests against chance with 95% confidence intervals are shown; p-values are FWER corrected (Bonferroni) for 2 comparisons (left and right hemisphere) and thresholded: *<0.05, **<0.01, ***<0.005.

This decoding approach handles confounds differently than the regression approach. The decoding models were trained to predict character identity but the analysis was performed on the estimates for the non-correct character, holding the within character effect on neural pattern similarity constant. To reduce effects of time (e.g., the narrative structure and drift), each model split was trained on random decision trials from across the task. The analysis also used the end-of-task places, when all the characters had been interacted with the same number of times; place location differences couldn’t reflect differences in familiarity. To rule out that the characters’ temporal proximity explained the effect (i.e., that participants’ interaction decisions are more similar the closer in the narrative they are, so that temporally close characters also have close social places at the end of the task), we performed the following control analysis. For each participant, we correlated (Pearson’s r) the characters’ pairwise end-of-game place distances with their average pairwise narrative-based temporal distances (the average time interval between different characters’ decision trials). The correlation coefficients were Fisher z-transformed and entered into a right-tailed 1-sample t-test: if temporal distance explains the relationship between neural decoding and place distance, then we expect a positive effect. The analysis was repeated for the squared temporal distances.

### 2D social angle replication

Two previous studies found that left hippocampal BOLD activity correlated with the angle of the vector from the participant’s theoretical point-of-view to the current character’s social place (Tavares et al., 2015; Zhang et al., 2022). This analysis assumes that while making decisions, participants represent the characters’ locations in an “egocentric” fashion: from their own first-person, subjective point-of-view. We sought to replicate this effect in this study’s validation sample (note: the initial sample in this study is from Tavares et al., 2015).

To do this, parametric modulation GLMs were fitted to each participant’s (smoothed) time series with the 2D social angles as a parametric regressor on the decision trials. The beta estimates for the parametric regressor were contrasted against the implicit baseline (i.e., unmodeled BOLD activity). To replicate the effect in the validation sample, we created an ROI from the results of the previous two studies. We re-ran the analysis in the initial sample and combined the resulting peak left hippocampal coordinate with the peak left hippocampal coordinate reported in Zhang et al. (2022) to generate a sample size weighted coordinate around which we defined a spherical ROI (radius = 5mm). This ROI was used to test the validation sample’s parametric modulator contrasts for significant positive effects against 0 with a 1-sample t-test. For more information see **Parametric modulator GLM** in the supplement/

### Long axis analyses

There may be a local to global representational gradient along the posterior to anterior hippocampus (i.e., long axis); if so, anterior hippocampal patterns should 1) be more correlated (Brunec et al., 2018) and 2) be better approximated by lower-dimensional embeddings than posterior patterns. To test the first prediction, on the participant level the anterior and posterior patterns were separately pairwise correlated (Pearson’s r), averaged and Fisher z-transformed. The anterior and posterior averages were then statistically compared against each other in ANCOVA, with follow up right-tailed paired t-tests. To test the second prediction, we first had to estimate the dimensionality of hippocampal patterns along the long axis. To do this, we applied multidimensional scaling (MDS) to the anterior and posterior hippocampal pattern correlation distances (1 – Pearson’s r) for each participant. MDS projects distances into lower-dimensional embeddings while maximally preserving the distance structure: smaller distances between the original distances and the projected (lower-dimensional) distances indicates a better fit (more details can be found in **Multidimensional scaling analysis details** in the supplement). We computed every n-dimensional anterior and posterior hippocampal projection in range [2, 10] and for each n compared the goodness-of-fit (Kruskal’s stress: a smaller value indicates a better fit) for the anterior and posterior with ANOVA and follow-up one-sided paired t-tests.

Control analyses were used to ensure that any BOLD pattern differences along the longitudinal axis were not due to differences in temporal signal-to-noise ratio (tSNR). tSNR estimates data quality across the BOLD time course, and is calculated as the time course’s mean divided by its standard deviation. For each participant, tSNR was calculated in the warped but unsmoothed functional images for all left and right anterior and posterior hippocampal voxels, and was averaged across each ROI. We used ANOVA to test whether tSNR differences in the anterior and posterior hippocampus related to pattern similarity or MDS fits. If there was a statistically significant relationship between tSNR and the outcome of interest, we included tSNR as a covariate in the main analysis.

## RESULTS

### Participants’ behavior is idiosyncratic

To identify a neural correlate of social place, the places should vary across participants; otherwise a place effect would be hard to disentangle from a shared confound across participants. To test whether places differ across participants, the angles from the neutral task origin (0,0) to each participant’s average affiliation and power location (i.e., the average of their +1/-1 decisions on the two dimensions) were tested for uniformity across 360 degrees with Rayleigh’s test for uniformity. The null hypothesis of a uniform distribution couldn’t be rejected (z=1.54, *p*=0.21): the average places span the entire 360 degree space (angle *M*=45.68, *SD*=73.57). Thus, there was no evidence for a common behavioral trajectory across the participants.

### Post-task character placements support 2D social space’s ecological validity

Do the social places we model from the participants’ decisions reflect the participants’ subjective perceptions of the characters’ social places? To test this, we compared the average distances from the task places to the post-task placements to the average distances between the task places and randomly generated locations. As expected, the post-task placements were closer to the end-of-task places (*t*(31)=-5.05, *p*<2e-5, 95% CI=[−0.266, −0.113]), indicating that the 2D representations we model from the task reflect the participants’ own perceptions. These results suggest our social space behavioral modeling has ecological validity.

### There is a hippocampal social place representation

#### Representational similarity regression shows social place effect in hippocampus

To test our main hypothesis that the hippocampus represents 2D social places, we first used ANOVA on the initial sample’s social place regression beta estimates. As expected, hippocampus pattern dissimilarity increased with the place distance: hippocampal betas were significantly different from 0 (average hippocampus effect: *F*(1,17)=11.378, *p*=0.004, pes=0.401), with no effect of hemisphere (*F*(1,17)=1.362, *p*=0.259). The place effect was in the predicted direction: farther apart places had more dissimilar hippocampal patterns (right tailed t-test for the averaged hippocampal betas: *t*(17)=3.373, *p*=0.0018, 95% CI=[0.0006, 0.0028], d=0.795). Further, the behavioral control models did not have significant hippocampal effects (all average effect *p*s>0.08; all hemisphere effect *p*s>0.2), suggesting this effect is strongest when both dimensions are considered together.

We wanted to ensure this hippocampal 2D social place effect was not spurious or specific to this sample by replicating it in the validation sample. Power analysis estimates ~97% power to detect an effect of the same size (d=0.795) in the validation sample (n=32) with a right-tailed 1-sample t-test at alpha=0.05, suggesting this is a statistically well-powered analysis: there is a very low probability of a false negative. The predicted social place effect replicated in the validation sample (average hippocampus effect: *F*(1,31)=6.142 *p*=0.019, pes=0.165; no effect of hemisphere: *F*(1,31)=0.252, *p*=0.619), with the same positive relationship between social place distance and hippocampal pattern dissimilarity (*t*(32)=2.478, *p=*0.009, 95% CI=[0.00045, 0.0047], d=0.438).

Our predictions were specific to the hippocampus: we hypothesized that a social place-like representation is encoded by hippocampal place cells; thus the hippocampus should best reflect these representations in its fMRI signals. To estimate the social place effect’s regional specificity, we performed control ANCOVAS with region and hemisphere as within-participant factor and sample as a between-participants factor. An ANCOVA with M1 had a significant effect of region (type II ANOVA *F*(1,48)=5.708, *p*=0.021, pes=0.106), but no effect of hemisphere (*p*=0.994) or sample (*p*=0.634). As expected, the social place beta estimates for the hippocampus were significantly larger than for M1 (*t*(49)=2.365, *p*=0.011, 95% CI=[0.0004, 0.0049], d=0.334); further, the M1 betas themselves were not statistically different from 0 (*p*=0.62). ANCOVA with PCG showed similar results: a significant effect of region (type II ANOVA *F*(1,48)=9.386, *p*=0.004, pes=0.164), with larger hippocampal betas (*t*(49)=3.001, p=0.002, 95% CI=[0.0011, 0.0059], d=0.424) and PCG betas that are not themselves greater than 0 (*p*=0.88). As such, with respect to the regions tested here, the social place effect is specific to the hippocampus.

#### Anterior hippocampus is significant in whole-brain analysis

To corroborate the hippocampal social place effect and explore the rest of the brain, we performed a whole-brain searchlight representational similarity regression. We expected that a place representation effect would be seen in the hippocampus and would be strongest in the anterior aspect given its possible role in representing abstract structure (Strange et al., 2014). Indeed, the voxels that survived multiple comparisons correction were located predominantly in the anterior left hippocampus (peak: −21/-15/-22; voxel extent=117 2.1mm^3^ voxels) **(fig. 4)**. A cluster in the right thalamus also survived (peak: 11/-9/8; voxel extent=143), reflecting its role in hippocampal place field stability (Cholvin et al., 2018)—perhaps as a mediator of medial prefrontal cortex to hippocampal interactions. These whole-brain results support the hypothesis that the hippocampus encodes a maplike representation of the characters’ changing social places across social interactions.

**Fig 4.**
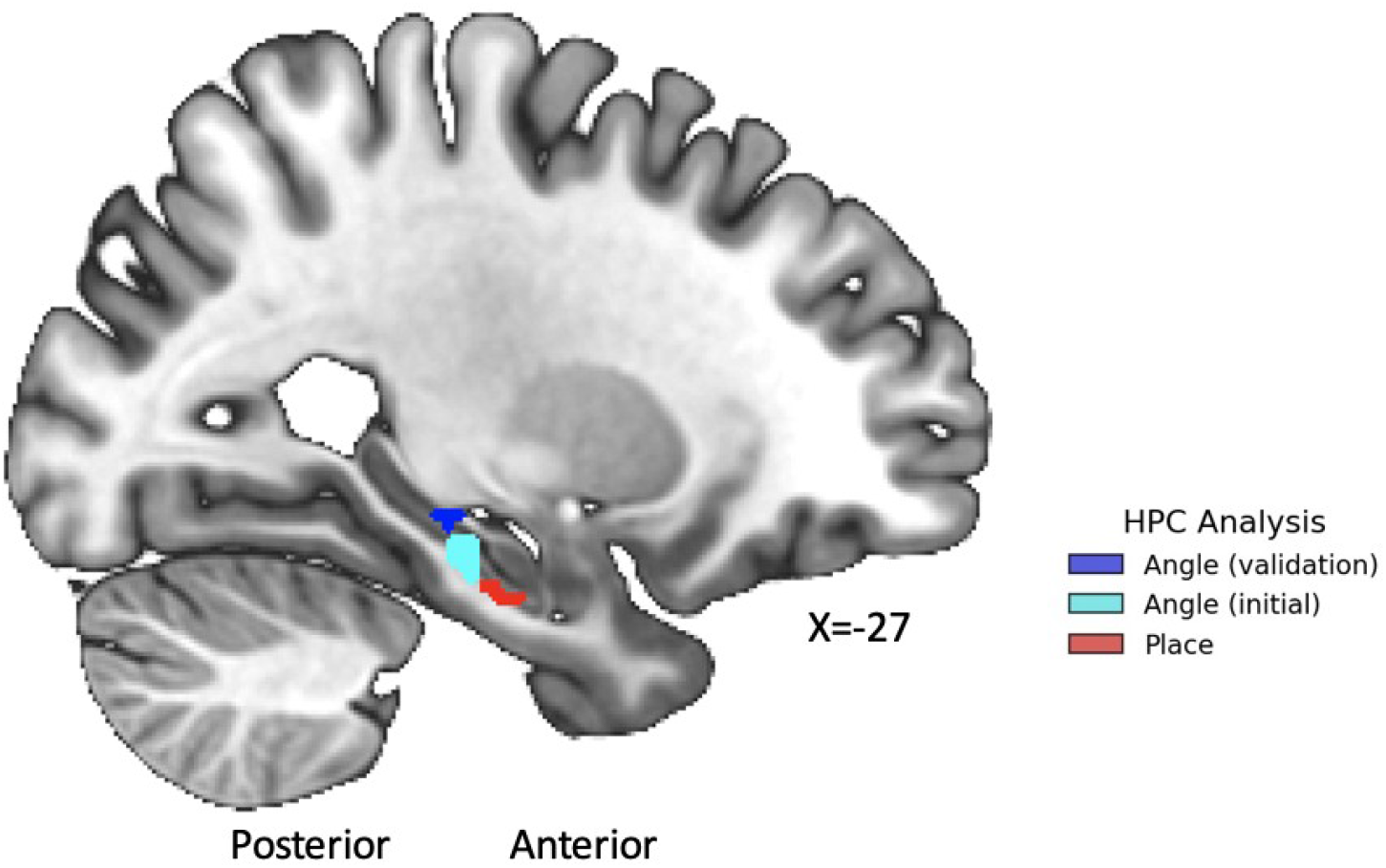
Place and angle clusters are differentially distributed along the long axis of the left hippocampus. The place cluster (red) is more anterior along the hippocampal long axis than the two angle clusters (blues). The clusters are binarized and shown on a slice (MNI X=-27) that can show all three clusters.

#### The hippocampal social place representation is related to post-task memory

If hippocampal social map-like representations during social interactions are important for subsequent social memory (Schafer and Schiller, 2018), then we may see a positive relationship between the post-task memory scores and hippocampal beta weights from the ROI representational similarity regression. To test this, we regressed (OLS) the post-task memory onto the average hippocampal betas for the predictor that tracked relationship changes: social place and familiarity. The relationship with memory was significant for both familiarity (*t*(31)=2.747, *P*_FDR_=0.01) and social place (*t*(31)=1.968, *P*_FDR_=0.029) (adj. R^2^=0.221). Thus, social memory may relate to hippocampal representations of 2D social places across interactions.

### Supporting evidence for hippocampal social place representation

#### Close social places have more similar hippocampal patterns

The hippocampal pattern similarity of trial pairs in 2D space corroborated the hippocampal social place effect: the average hippocampal pattern similarity (Pearson’s r) varied as a function of the place distance (χ^2^(2)=13.637, *p*=0.0011). The pattern was consistent across the three bins of place distances: the closest trials had the highest average pattern similarity (pattern similarity *M*=0.0284), followed by the next closest trials (*M*=0.0268) and then the farthest trial pairs (*M*=0.025).

There was no such relationship for M1 (*p*=0.58) or PCG (*p=*0.097); the average difference between the closest and farthest trials was also greater in the hippocampus than in both control regions (HPC>M1: *t*(49)=1.739, *p*=0.044, 95% CI=[-0.0007, 0.0097], d=0.25; HPC>PCG: *t*(49)=2.198, *p*=0.016, 95% CI=[0.0005, 0.011], d=0.31). Importantly, only trial pairs with different characters were included in these analyses, to control for the higher pattern similarity associated with same character trial pairs. Yet, even under these minimal assumptions we see a statistically significant hippocampal social place effect.

#### Hippocampal decoders confuse characters who are nearby in social space

We also used a probabilistic decoding approach to provide convergent evidence for hippocampal social place mapping. If hippocampal patterns represent the characters as places in 2D social space, a character decoder trained only on those patterns may confuse characters with nearby social places more often.

We first decoded character identity from the hippocampus patterns (average hippocampal accuracy: Initial *M*=35.6%, *SD*=6.2%, Validation *M*=38.4%, *SD*=5.2%) with above chance accuracy across samples and hemispheres (average hippocampal effect: type II ANOVA *F*(1,48)=493.841, *p*<6.6e-27, pes=0.911; *t*(49)=21.767, *p*<3.9e-27, 95% CI=[35.8%, 38.9%], d=3.08). The hippocampal decoding accuracies also strongly correlated with the averaged character identity betas in the hippocampus from the representational similarity regression (OLS, controlling for sample: *t*(47)=3.572, *p*=0.00083, 95% CI=[1.297, 4.641], adj. R^2^=0.261), suggesting the decoding reflects character identity and not just task, behavior or time-related confounds, which are well controlled for in the regression analyses. Decoding in the hippocampus was also more accurate than in M1 (paired t-test: *t*(49)=4.947, *p*<4.7e-06, 95% CI=[2.9%, 6.9%], d=0.6996), suggesting that hippocampal patterns contain more (linearly decodable) character information than M1. In contrast, PCG patterns provided comparable character decoding as hippocampus (paired t-test: *p*=0.36).

We then tested whether the characters’ end-of-task 2D social places relate to hippocampal decoder confusability. As predicted, the average hippocampal confusability effect was greater than chance (type III ANOVA *F*(1,48)=8.761, *p*=0.005, pes=0.154; *t*(49)=3.185, *p*=0.00125, 95% CI=[50.857%, 53.78%], d=0.45). There were no main effects of the sample or hemisphere (both *p*s>0.2) but there was a significant interaction between sample and hemisphere (type III ANOVA *F*(1,48)=6.851, *p*=0.012, pes=0.125): the initial sample had a larger effect in the right than left hippocampus (right *M*=54%, left *M*=50.2%).

The hippocampal confusability effect was greater than the confusability effect in M1 (t(49)=2.664, *p*=0.005, 95% CI=[0.6%, 4.35%], d=0.376) and PCG (*t*(49)=1.689, *p*=0.048, 95% CI=[-0.3%, 3.49%], d=0.24) (see **fig. 3**). That the hippocampal character confusability effect is greater than the PCG effect, despite comparable character decoding, suggests that the confusability effect is not just a function of decodable character-related information but instead reflects social place-related information. The control models all also had insignificant hippocampal confusability effects (all *p*s>0.05), suggesting that social place was a better explanation for these data. Closeness in time also doesn’t explain decoder confusability: the characters’ end-of-task place distances were not correlated with their temporal distances (*p*=0.229; distances^2^ *p*=0.223), suggesting the effect can’t easily be explained by time-related variables, such as the characters’ co-occurrence in scenes or temporal autocorrelation of the BOLD signal. As such, the decoders’ confusion about character identity based on latent social places supports the effect seen in our regression analyses: hippocampus patterns contain information about the characters’ 2D social places.

### Egocentric social angle effect replicates in left hippocampus

In the validation sample, we replicated the left hippocampal effect of the trial-wise social angle to the character from the theoretical point-of-view of the participant. To first generate an ROI to test for this effect in the validation sample, we combined the peak left hippocampus voxel from a re-analysis of the initial sample (peak: −24/-22/-22; *t*(17)=5.38, *p*_FWER_=0.011, with a *p*<0.05 cluster extent=18 3mm^3^ voxels) and the peak left hippocampus voxel from a recently published replication (Zhang et al., 2022) to calculate a sample size weighted average peak coordinate (−23/-26/-11), around which we defined a spherical ROI. Then, using this ROI, we replicated the angle parametric modulation effect in the validation sample (peak: −25/-24/-7; *t*(31)=3.02, *p*_FWER_=0.026, *p*<0.05 cluster extent=18 2.1mm^3^ voxels).

### Social angle and social place are complementary effects

The angle analyses’ hippocampal clusters, as well a cluster from a recent replication (Zhang et al., 2022, were more posterior than the place searchlight’s cluster (see **fig. 4**). This may relate to how locations are compared in these analyses. In the place analysis, Euclidean distance is used: a small Euclidean distance indicates that places are nearby in space, irrespective of orientation. In the angle analysis, cosine similarity is used: a high cosine similarity indicates that places are in the same general direction from the point-of-view of the participant, irrespective of distance. These differences mirror place and angle differences seen in physical space: in a recent fMRI study, the participant’s location was correlated with anterior hippocampal patterns while the participant’s heading direction was correlated with posterior patterns (Kim et al., 2017).

The two analyses also modeled BOLD events differently, reflecting different assumptions about the underlying neuronal events. The place pattern analyses modeled the BOLD response at the onset of the decision trial, assuming that the updated location would be encoded in neural activity at the beginning of the decision points. The parametric modulation analysis modeled the BOLD event for the duration of the reaction time, with the assumption that representing angle from the participant’s point-of-view to the updated location would unfold over the course of the decision deliberation period. This raises a question, to be answered in future studies: are places and angles differentially represented (e.g., patterns versus magnitudes) as a function of time-related behavioral demands (e.g., where in the decision-making process the participant is)? The analyses also differed in multiple other ways, (e.g., multivariate versus univariate; different smoothing kernels) thus caution should be taken in interpreting differences between the results.

### Anterior and posterior hippocampus have different functional properties

If there is a representational gradient for spatial scale along the hippocampus’s longitudinal axis with the posterior representing using ‘finer’ activity patterns to represent details and the anterior using ‘coarser’ activity patterns to represent larger structures (Brunec et al., 2018), then there should be greater average decision trial pattern similarity in the anterior hippocampus than the posterior. We tested this with ANOVA on the hippocampal pattern correlations between all trials for the anterior versus the posterior. As expected, there was a significant effect of hippocampus segment (type II ANOVA *F*(1,48)=4.569, *p*=0.038, pes=0.087) with higher pattern similarity in the anterior than posterior hippocampus (*t*(49)=2.146, *p*=0.018, 95% CI=[0.00039, 0.0119], d=0.303). Temporal SNR differences in the anterior and posterior did not significantly relate to pattern similarity (*p*=0.09), suggesting this long axis difference is driven by task demands: greater anterior hippocampal pattern similarity might reflect a low-dimensional representation of 2D social space (**fig. 5A**).

**Fig 5.**
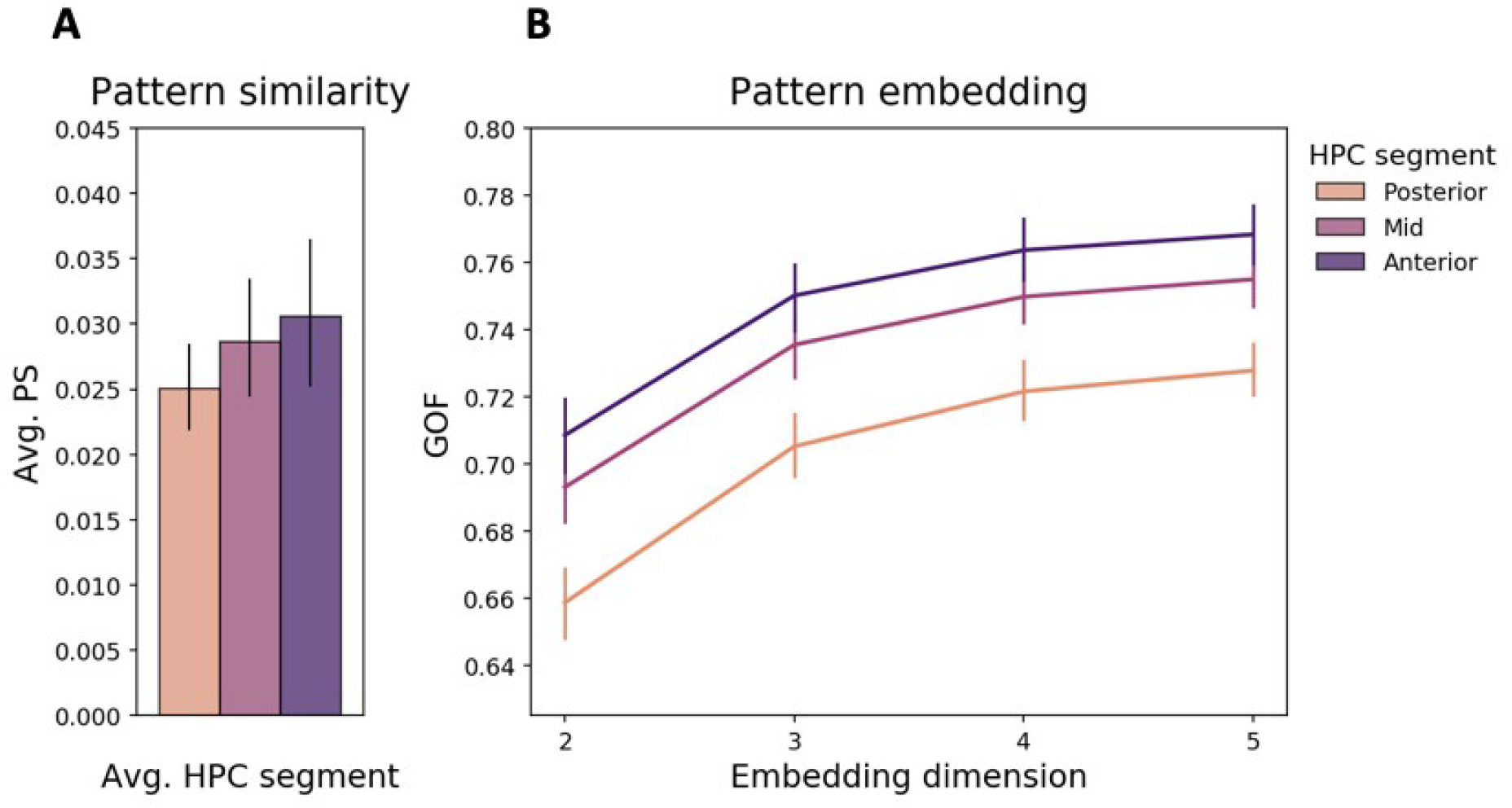
Patterns in the anterior hippocampus are more correlated and lower dimensional than posterior patterns. (A) The anterior hippocampus has higher average pattern similarity across the decision trials than the posterior hippocampus. The mid hippocampus is shown as well to demonstrate the gradient-like pattern: pattern similarity may gradually increase along the long axis. (B) The goodness of fit metric is 1 – Kruskal’s stress; larger values indicate a better fit. Low dimensional embeddings explained the correlation distances between anterior hippocampus patterns during decision trials better than patterns from posterior hippocampus.

If this representational gradient exists along the hippocampal long axis, anterior patterns might be better approximated by low-dimensional embeddings than posterior patterns. To test this, we used multidimensional scaling (MDS) to project the anterior and posterior pattern correlation distances (1 – Pearson’s r) into lower dimensional spaces, and then tested if the goodness of fit of these projections varied along the long axis. As expected, pattern distances in the anterior hippocampus were better approximated (i.e., had better fits) by low-dimensional embeddings than the distances in the posterior hippocampus, for all dimensions tested (dimensions: [2,10]; Likelihood Ratio Test *p*s<0.0002). These models aso included tSNR, since tSNR and MDS fits were statistically related (*p*<2e-15); the robustness of this dimensionality effect suggests that the relationship between the long axis and dimensionality is intrinsic to the BOLD signal and not noise (**fig. 5B**).

## DISCUSSION

### The hippocampus represents latent abstract places

We showed that a hippocampal latent place-like representation is present during naturalistic social interactions, suggesting that the hippocampal place code does more than represent locations in physical space: its function is domain general. We used two independent samples and four different analyses: an ROI and searchlight representational similarity analysis, a decoding probability analysis and an average pattern similarity analysis. Hippocampal place effects were seen both with *a priori* ROI and whole-brain searchlight analyses. The interpretation that our findings reflected latent, 2D social places was bolstered by the controls included in the regression models: these patterns were not explainable by reaction time or button press, time-related drift, narrative structure, character identity, or the number of interactions with a given character. Our confidence in the effect is strengthened by a character decoding confusability analysis and a social place average pattern similarity analysis. All these results were validated in the second sample and compared against control regions, suggesting these effects are stable and relatively specific to the hippocampus. Further, the two social dimensions (affiliation and power) were not perceivable during the game, and there were no instructions about their presence or reinforcement of the locations. Participants merely made decisions in the interactions as they would in the real-world—suggesting these latent place representations emerged entirely from the dynamic social interactions.

The decoding confusability analysis further demonstrated that hippocampal patterns contained information about characters’ latent 2D social places. These results also suggest that the characters’ identities and their social places may be represented conjunctively, similar to hippocampal conjunctive representations of object identity and location (Manns and Eichenbaum, 2009). The hippocampus may represent both *who* we are interacting with and *where* they are in our social space (“who is where”).

### Implications for social mapping

The whole-brain searchlight analysis revealed an especially strong left anterior hippocampus effect, following other fMRI studies finding anterior place-like representations in both physical (Morgan, LK, MacEvoy, SP, Aguirre, GK, Epstein, 2011) and conceptual spaces (Theves et al., 2019), and across 189 “place”-related fMRI studies (automated meta-analysis: Yarkoni et al., 2011). The anteriorness of this effect also suggests a connection to other work that posits a posterior-to-anterior local-to-global representational gradient along the long axis of the hippocampus (Brunec et al., 2018; Strange et al., 2014). The anterior hippocampus’ relatively larger place fields and denser place coding could cause greater ensemble-level similarity relative to the posterior: indeed, in our data average pattern similarity during the decision trials was higher in the anterior than the posterior hippocampus. This property could abstract the low-dimensional structure from (high-dimensional) experience: consistent with this, anterior hippocampal patterns were better approximated by lower-dimensional embeddings than posterior patterns. Other research suggests social memory in particular may relate to this anterior and posterior distinction (Rao et al., 2019). These results are all consistent with the anterior hippocampus representing global information, like the places on a map.

Flexible navigation between places may also require vector computations. Place cells become directionally tuned as behavior gains salience (Markus et al., 1995), suggesting navigational demands engage conjunctive place and direction representations. The posterior hippocampus in particular seems to encode directional vectors from the animal’s current location to goal locations (Sarel et al., 2017). Similarly, as participants make their decisions in our social navigation task, the left mid/posterior hippocampus parametrically tracks the directional angle from the participant’s theoretical point-of-view to the updated social places of the characters (Tavares et al., 2015; Zhang et al., 2022)—a finding we replicated here in our validation sample. Previous fMRI work in physical space mirrors this long-axis distinction: the anterior hippocampus represents locations and their global properties, whereas the posterior hippocampus represents directions and detailed trajectories (Howard et al., 2014; Javadi et al., 2017; Kim et al., 2017). As such, mapping and navigation may engage both anterior signals related to global place representations and posterior signals related to vector-based navigation. This view could help make sense of the thalamic cluster seen in the place searchlight analysis: thalamus helps stabilize posterior hippocampal place fields (Cholvin et al., 2018), perhaps by mediating a medial prefrontal cortex to hippocampal connection that serves to differentiate similar spatial representations: for example, between similar spatial locations and trajectories. This suggests an intriguing framework to understand navigation in abstract social space: at decision points, the anterior hippocampus may represent global place information (“where am I?”, “where should I be?”) while the posterior hippocampus represents navigational vectors between places (“what direction should I take?”), possibly to guide navigational action selection (“I will go that way”). These ideas require rigorous testing in future work.

Our results also complement other work showing social mapping-like effects. In learning tasks with social information, the hippocampus tracks distances in 1D space (Kumaran et al., 2016), as well as inferences about 1D (Kumaran et al., 2012) and 2D social spaces (Park et al., 2020). These studies did not show place effects, and they had learning phases for participants to explicitly learn the relevant social dimensions (i.e., not latent), often using pairwise comparisons between characters (i.e., not naturalistic). Moreover, the participants were observers of, and not navigators of, the social spaces: there was no true navigation-like behavior. Thus, while previous studies have shown evidence for hippocampal computations of relationships between locations (e.g., directions, distances and inferences between places), this is the first study to provide direct evidence for the latent social place representations themselves.

### fMRI can detect place-like representations

But how could fMRI BOLD signal reflect the activity of place cells? If place cells with nearby firing fields are non-uniformly distributed in the hippocampus (i.e., if they cluster), then we may be able to detect a place-like pattern across fMRI voxels. Rodent work has suggested there may be some anatomical structure to place cell organization, whereby hippocampal place cells with nearby place fields cluster (Dombeck et al., 2010; Hampson et al., 1996, 1999). Most importantly, multiple previous fMRI studies have shown that physical locations can be decoded from hippocampal BOLD patterns (Hassabis et al., 2009; Kim et al., 2017; Sun et al., 2021) and that the pattern similarity decreases with the distances between locations, as is expected from a map-like place code (Morgan, LK, MacEvoy, SP, Aguirre, GK, Epstein, 2011). As such, place cell or ensemble activity may be detectable at the fMRI voxel level as place-like signals—even latent place-like signals.

### Future directions and conclusion

Future work should probe hippocampal social place-like representations more closely: do they reflect social inferences and/or predictions (Stachenfeld et al., 2018)? How do they interface with other space-like representations in the medial temporal lobe, like grid-like representations, or decisionmaking mechanisms in the frontal lobes? Other work can also probe for similar processes in other species: for example, non-human primates use conspecifics’ relative locations on affiliation and power dimensions to make inferences and manage conflict (Cheney and Seyfarth, 1990), while their own locations on these dimensions relate to their reproductive success (Feldblum et al., 2021). Circuit manipulations to alter social representations in the hippocampus (e.g., in anterior CA1: Rao et al., 2019) could alter inferences on these dimensions and downstream behaviors (e.g., both cooperative and competitive), offering a view onto the functional relevance of these representations. Crossspecies variation in latent social structures and how they are navigated could also provide clues about the roles of genetics and learning in abstract social representations. There may be clinical relevance to these processes as well: many disorders feature both hippocampal abnormalities and social dysfunction consistent with mapping deficits (Schafer and Schiller, 2019).

In conclusion, we provide evidence for a domain general hippocampal place-like representation during a social interaction game. The social dimensions were devoid of perceivable features, and the participants were not cued about the underlying dimensions or reinforced to learn specific character locations. Further, the presence of the map was inferred from brain patterns during naturalistic behavior, where there was no explicit task requirement to express the characters like locations in a map. Thus the place-like representations seen here emerge from participants’ naturalistic interaction decisions and constitute a fully domain general and highly abstract map-like coordinate system. These results clear a path for furthering the study of hippocampal cognitive mapping in all domains of experience.

