## Supplemental Info for "Hippocampal Place-like Signal in Latent Space"

**This PDF file includes:**

Materials and Methods

**Social Navigation Task details**

***Counterbalancing***

The assignment of the button response box keys to choice direction (i.e., +/-) was counterbalanced. The gender presentation of the characters was also counterbalanced within the task such that half of the characters presented as men and presented as women. The assignment of gender to specific characters was further counterbalanced across participants: i.e., half of the participants received a narrative version where the gender of specific characters was flipped from the other half. Further in the validation sample, the skin color of the characters was counterbalanced in the same way as gender, with darker and lighter skinned versions of the same underlying character image. The version of the task for the initial sample had the same text and options but more cartoon-like character images (Tavares et al., 2015).

***Participant's theoretical point-of-view***

For several analyses (e.g., angle parametric modulation, post-task placement) a theoretical point-of-view of the participant was assumed: i.e., an abstract first person point-of-view for the participant to the changes in their social space. We modeled this point-of-view as the maximum affiliation value a character can have at the end of the task (+6) and the neutral value on the power dimension (0), for a point-of-view location of (+6, 0). Thus, the characters can move up or down on the power axis relative to the participant and gain as much affiliation with the participant as the participant's affiliation location allows (+6).

**Behavioral analyses**

***Affiliation and power dimensions are orthogonal in behavior***

We model each participant's social space as Euclidean, which assumes that the affiliation and power dimensions are orthogonal to one another and can be represented together as a Euclidean plane, allowing us to use the Euclidean distance function as our metric. We wrote the narrative so that choosing either option in a given decision was perceived as changing the relationship with the character mainly along either the affiliation or power dimensions, approximately the same magnitude but with opposite signs (directions). In other words, in a power (or affiliation) decision, the two options both would mainly change the power (or affiliation) dynamics in the current relationship, with one choice increasing the participant's power and the other increasing the character's power (or affiliation), both in about equal amounts.

We validated this assumption empirically. To do so, we estimated the orthogonality of the affiliation and power decisions and the orthogonality of the locations in the post-task social space placement task. If the dimensions are orthogonal, they should be uncorrelated: behavior (or post-task placement for the validation sample) on one dimension should not predict the other. To test this, each participant's character-wise affiliation and power values (5 pairs per participant) were correlated (Pearson's r); the coefficients were then Fisher z-transformed and tested against 0 with a 1-sample t-test. The null hypothesis (of 0 correlation) could not be rejected for the task average locations ($p$=0.998) or for the post-task placement ($p$=0.83). Affiliation and power decisions were also uncorrelated across participants (task average: r=0.224, $p$=0.12; post-task placement: $p$=0.69). These complementary results, in both the task behavior and post-task placements, support the hypothesis that affiliation and power decisions are orthogonal in this task.

**fMRI details**

#### *Image acquisition*

The task was collected in a single imaging run of approximately 26 minutes. Imaging parameters differed between the two samples, so each is described separately.

- Original sample: T2*-weighted functional images were collected on a Siemens Allegra 3T scanner, with a single-shot EPI pulse sequence with the following parameters: flip angle = 90 deg, TE = 35 ms, TR = 2,000 ms, 36 slices, 64 x 64 matrix, voxel size = 3 $mm^3$. T1-weighted images were obtained with an MPRAGE protocol with a voxel size = 1 $mm^3$.
- 
- Validation sample: T2*-weighted functional images were collected on a Siemens Skyra 3-Tesla (3T) scanner (Siemens, Erlangen, Germany), with a multiband slice echo-planar imaging (EPI) pulse sequence with the following parameters: acceleration factor = 7, flip angle = 60 deg, echo time (TE) = 35 ms, repetition time (TR) = 1,000 ms, 70 slices, 108 x 108 matrix, voxel size = 2.1 $mm^3$. T1-weighted images were obtained with a magnetization-prepared rapid gradient-echo (MPRAGE) protocol with a voxel size = 0.8 $mm^3$.

#### *Image preprocessing*

The same preprocessing steps were performed for all participants using SPM12 (Wellcome Trust Centre for Neuroimaging). To correct for participant motion, functional images were re-aligned to the first volume using a standard six parameter rigid body transformation and unwarped to account for magnetic field inhomogeneities. The realigned images were slice-time corrected to the middle slice, then co-registered in alignment with the MPRAGE to the mean unwarped image. The MPRAGE image was then segmented into 6 tissue classes and the resulting forward deformation parameters were used to normalize the unwarped, coregistered functional images to the standard MNI (Montreal Neurological Institute) template. Images were resliced to the same voxel size as the respective dataset's functional images. The unsmoothed time series images were used in the general linear models (described below) to preserve the multi-voxel patterns.

#### *Regions of interest (ROI) definitions*

ROI analyses were performed to test our 2D social place predictions in the hippocampus. Probabilistic right and left hippocampus masks were defined from the Harvard-Oxford Atlas in FSL (FMRIB Software Library), co-registered and re-sampled to the validation functional image dimensions, and binarized at 10% probability. We also divided the hippocampus into anterior and posterior portions, in approximately equal fractions of the long axis of the hippocampus: anterior from Y=[-18,-4] and posterior from Y=[-40,-30] (following Collin et al., 2015). Primary motor cortex and precentral gyrus masks from the same atlas were thresholded at 25% probability and served as control regions.

As distributed patterns are likely closer to the underlying neuronal processes (e.g., information encoded in spatially distributed neuronal ensembles) than activity magnitudes of single voxels, we used multivoxel pattern analyses to test our hypotheses. Multivoxel analyses can reveal spatially distributed representations that are undetectable to mass univariate methods that assume each voxel has an independent activity profile. For a given ROI, each participant's voxelwise, unsmoothed beta weights were extracted using the binarized ROI mask on the trial-level GLM beta maps. Any voxels that did not exceed 50% of the global signal were excluded prior to statistical analysis.

***General linear modeling (GLM)***

GLMs for both the pattern analyses and parametric modulation analyses were fitted in SPM12. For all GLMs, the microtime resolution was set to the number of slices collected and the microtime onset was set to the middle slice. The six motion parameters (3 translation and 3 rotation) from preprocessing realignment were included in the design matrices (without hemodynamic response function convolution) to regress out residual motion-related variance. To reduce autocorrelation in the time series, a high-pass filter of 128 seconds was used to remove low frequency signals and prewhitening was performed with SPM's FAST algorithm. Beta weights were estimated for every voxel whose average signal exceeded 50% of the global signal.

***Trial-wise GLM***

Trial-wise GLMs were fitted to each participant's unsmoothed time series. The design matrix contained separate condition regressors for each decision trial onset. The narrative trial (non-decision trials, where the characters are speaking or background information is provided) onsets were modeled by an additional condition regressor. The blood oxygenation level dependent (BOLD) signal associated with each regressor was modeled at the start of each decision trial by a stick function convolved with a canonical hemodynamic response function. The resulting beta images for the initial sample were then resampled to match the functional images from the validation sample (which has higher spatial resolution), so that the samples are more directly comparable in subsequent analyses.

***Parametric modulator GLM***

Parametric modulator GLMs were fitted to each participant's smoothed (6mm at FWHM) time series, following Tavares et al., 2015. The design matrix included separate regressors for the narrative trials and decision trials. A parametric weight was applied to the decision trials based on the 2D angles. To calculate these angles, we defined a directional vector from a theoretical point-of-view of the participant (max affiliation, neutral power: 6, 0) to each trial's 2D place and another vector from the participant's point-of-view to the max affiliation and power coordinate (6, 6). A third axis was also included to control for the number of previous interactions with the given character (1-12; i.e., familiarity with the character). The cosine of the angles between these (trial-wise) vectors were entered into the design matrix as parametric modulators of the decision trials. The cosine of the angle between any two vectors *u* and *v* is given by the dot product of the vectors divided by the product of their norms (i.e., the cosine similarity between the vectors):

$$\cos(\text{angle}) = \text{dot}(u, v) / (\text{norm}(u) * \text{norm}(v))$$

All task predictors (narrative, decisions and angles) were modeled with a boxcar stimulus convolved with a canonical hemodynamic response function. The decision and angle regressors events were modeled from the start of the decision trial to the length of the reaction time; the narrative trials as the full duration of the trial.

### Representational similarity regression details

#### *Social navigation distance matrices*

Several different models of participant's behavior ("social navigation") were tested (described in **Representational similarity regression analyses** in the main text) in their ability to explain the data. All models compared the decisions to each other using Euclidean distance (2D or the 1D absolute value difference); the models differed in how they represented each decision trial. Each model's trial-level representation is described below:

2D place: location on the affiliation and power dimensions (i.e., cumulative sum of affiliation and power decisions)

1D affiliation/power place: location on the affiliation or power dimension only

1D place: location on the currently active dimension

1D positive place: location on a negative-to-positive dimension, defined as the cumulative sum of the directional decisions (+1/-1)

2D distance from origin: vector length from the origin (0, 0) to the affiliation and power location

2D distance from POV: vector length from the theoretical participant point-of-view (6, 0) to the affiliation and power location (see ***Participant's theoretical point-of-view***)

#### *Control distance matrices*

Several other kinds of distance matrices were included in all representational similarity analysis (RSA) models as covariates of no interest. Each distance matrix of no interest is briefly described below.

Slide: the absolute value difference between the slide numbers (i.e., all trials, including narrative based trials and decision trials) of the compared trials.

Scene: the absolute difference between the scene numbers (the count of discrete scene changes) of the compared trials, to account for the scene based structure of the narrative (range=[0, 11]). Scene changes were defined at discontinuities in the narrative: i.e., a discrete change in physical place or time.

Character identity: categorical distance matrices for each of the five experimental characters capturing whether trial pairs were both from the given character (1) or at least one trial was a different (0) character.

Character familiarity: the absolute difference between the character-specific decision counts for the trial pairs being compared. (e.g., if one trial was the 11th decision with a given character and the other was the 3rd decision with a given character, the absolute difference value equals 8)

Button press: categorical distance matrix where trial pairs with the same button presses were coded as 1 and trial pairs with different button presses were 0.

Reaction time: the absolute pairwise differences between trial reaction times.

Temporal drift: cubic polynomial expansion of the onset intervals between trial pairs. fMRI pattern drift is primarily time dependent and can increase with the temporal proximity of events; as such, this should be accounted for in representational similarity analysis. We accounted for temporal drift patterns by regressing out their estimated effects (Alink et al., 2015). To estimate a temporal drift model that accounts for drift-related noise, we modeled hippocampal pattern dissimilarities as a function of trial onset interval models with polynomial expansions of increasing degrees, starting with polynomial order 1 (1=linear, 2=quadratic, 3=cubic, etc; we used a Huber estimator). To measure how much each successive polynomial term improved model fit, we calculated the difference in average adjusted $R^2$ from the previous model. The polynomial model where the adjusted $R^2$ decreased was where additional temporal drift covariates no longer helped capture drift-related variance in the hippocampal pattern dissimilarities. This point was reached at polynomial order 4, for both samples and in both the left and the right hippocampus; thus a polynomial order 3 model (i.e., a cubic expansion of the time differences) was included:

$$\beta_1 D_{Temporal}^1 + \beta_2 D_{Temporal}^2 + \beta_3 D_{Temporal}^3$$

For both ROI and searchlight approaches (details below), robust multiple regression was used to estimate the effect of 2D social places (and control variables) on neural pattern similarity.

1) Predictors: In addition to the target predictor (2D place distance, or one of the control model predictors), we included covariates of no interest (described above in ***Control distance matrices***), which were essential to our subsequent inferences: nearby trials are likely to have correlated activity patterns because of slowly varying factors, including scanner drift, attention and the narrative structure (e.g., character identity or discrete scene changes). As such, regressors were included to control for the neural pattern variance associated with temporal drift, slide and scene changes, character identities and familiarity, button press and reaction time. The upper triangles of the symmetric predictor matrices (excluding the diagonal) were flattened into 1-dimensional arrays. Continuous predictors were z-scored (mean centered and divided by their standard deviation); button press and character identity predictors were left as categorical (0 or 1). The z-scoring of the behavioral modeling distances (e.g., 2D place distance) helps make participants' beta estimates more directly comparable, given differences in how much "space" they covered with their decisions: the resulting beta values estimate the change in the neural pattern correlation distances with a 1 standard deviation increase in Euclidean distance.

2) Neural patterns: Each participant's pairwise neural patterns were correlated (Pearson's r), with the resulting correlations Fisher Z-transformed (to transform the coefficients from a bounded distribution of [-1, +1] to a normal distribution) and subtracted from 1 (to convert from a similarity into a dissimilarity metric: correlation distance).

These participant-level regressions were computed:

$$D_{NeuralI} = \beta_0 + \beta_1 D_{Place} + \beta_2 D_{Slide} + \beta_3 D_{Scene} + \beta_{4:8} D_{CharacterID1:5} + \beta_9 D_{CharacterFamiliarity} + \beta_{10} D_{BP} + \beta_{11} D_{RT} + \beta_{12} D_{Temporal}^1 + \beta_{13} D_{Temporal}^2 + \beta_{14} D_{Temporal}^3 + \varepsilon$$

3) Regression: A Huber estimator (alpha=0.001, epsilon=1.75) was used to regress neural pattern dissimilarity/distance onto the dissimilarity/distance predictors. Given we used betas from single trials (e.g., and not conditions), we wanted to reduce the chance of noise driving spurious results; Huber regression was used because it is robust to violations of OLS assumptions, especially the presence of outliers (i.e., "robust regression"). OLS minimizes the squares of the fitted residuals, which means large residuals have an exponential impact: outlier observations have a large influence on the best-fit line. In contrast, the Huber loss function minimizes the absolute loss for outlier observations (the loss function is linear for large residuals) and the squared loss for other observations (the loss function is quadratic for small residuals). Iterative optimization is needed to fit the line: we used the L-BFGS-B algorithm from scikit-learn. We also fitted the models using OLS to compare: this estimator produced highly similar results to Huber regression.

Note that the social navigation behavior distances (e.g., 2D place distance or other control distances) were z-scored; the beta estimates should be interpreted as the change in neural pattern correlation distances for a 1 standard deviation increase in the Euclidean distance between the trials, with the trials measured in the given way (e.g., as 2D places).

4) Statistical inference: To test for group-level statistical significance, the beta estimates for the target regressor (i.e., representing the participants' decisions: the 2d place or control models) for all participants were entered into ANOVA. For statistically significant effects ($p<0.05$), follow-up t-tests were used to establish the effects' directions. Right tailed t-tests for predicted relationships (and control comparisons) were used because our predictions were directional: e.g., our main hypothesis predicted that neural pattern distances and 2D place distances have a positive relationship. For our predictions, p-values are left uncorrected and instead are directly replicated in the validation sample. For more exploratory analyses (e.g., searchlight analyses), p-values are corrected for multiple comparisons, with subscripts indicating the error rate control method (FWER for family-wise error rate or FDR for false discovery rate).

### Multidimensional scaling (MDS) analysis details

MDS takes a distance matrix and projects the distances into a lower-dimensional space, preserving the pairwise distances between observations as well as possible. MDS models are fitted by minimizing a "raw stress" loss function: the SMACOF algorithm minimizes the sum of the squared distances between projected distances and original distances. The raw stress values were converted to a normed stress: Kruskal's stress (square root of raw stress divided by the sum of squared original distances), which is bound between 0 and 1. Smaller Kurskal's stress values indicate a better fit.

### Decoding confusability analysis details

To decode trial-wise character identity, multinomial L2 (i.e., "ridge") logistic regression was used (with lbfgs optimization and inverse regularization strength C=10). Logistic regression models were fitted to each participant's multi-voxel trial-wise neural patterns and used to estimate the probabilities for the five characters on each trial. Given the relatively large number of voxels (features) to trials (observations), a L2 norm penalty term was added to the logistic loss function, to penalize the squares of the parameter weights. This penalization retains all of the voxels as features but forces some of their estimated weights to be small. Six-fold stratified cross-validation was used, so that in each fold the training was performed on ten random decision trials from each (non-neutral) character (50 total; 83.3% of trials), which was then tested on the two remaining decision trials from each (non-neutral) character (10 total; 16.7% of trials). Excluding the three trials from the neutral character (given that they do no not have the same number of trials or a meaningful end-of-game location), this produced a 60x5 (trial x character probability) matrix per participant. For each trial, the maximum probability is the classification. Since the decoding models were trained on trials from across the whole task, we used the end-of-task locations, which reflect the participant's decisions across the whole task.
